# Plasmid analysis of NDM metallo-β-lactamase-producing Enterobacterales isolated in Vietnam

**DOI:** 10.1101/2020.03.18.996710

**Authors:** Aki Hirabayashi, Koji Yahara, Satomi Mitsuhashi, So Nakagawa, Tadashi Imanishi, Van Thi Thu Ha, An Van Nguyen, Son Thai Nguyen, Keigo Shibayama, Masato Suzuki

**Affiliations:** AMR Research Center, National Institute of Infectious Diseases, Tokyo, Japan; Department of Molecular Life Science, Tokai University School of Medicine, Kanagawa, Japan; Microbiology Department, Hospital 103, Military Medical University, Hanoi, Vietnam; Department of Bacteriology II, National Institute of Infectious Diseases, Tokyo, Japan

## Abstract

Carbapenem-resistant Enterobacterales (CRE) represent a serious threat to public health due to the lack of treatment and high mortality. The rate of antimicrobial resistance of Enterobacterales isolates to major antimicrobials, including carbapenems, is much higher in Vietnam than in Western countries, but the reasons remain unknown due to the lack of genomic epidemiology research. A previous study suggested that carbapenem resistance genes, such as the carbapenemase gene *bla*_NDM_, spread via plasmids among Enterobacterales in Vietnam. In this study, we characterized *bla*_NDM_-carrying plasmids in Enterobacterales isolated in Vietnam, and identified several possible cases of horizontal transfer of plasmids both within and among species of bacteria. Twenty-five carbapenem-nonsusceptible isolates from a medical institution in Hanoi were sequenced on Illumina short-read sequencers, and 13 *bla*_NDM_-positive isolates, including isolates of *Klebsiella pneumoniae*, *Escherichia coli*, *Citrobacter freundii*, *Morganella morganii*, and *Proteus mirabilis*, were further sequenced on an Oxford Nanopore Technologies long-read sequencer to obtain complete plasmid sequences. Almost identical 73 kb IncFII(pSE11)::IncN hybrid plasmids carrying *bla*_NDM-1_ were found in a *P. mirabilis* isolate and an *M. morganii* isolate. A 112 kb IncFII(pRSB107)::IncN hybrid plasmid carrying *bla*_NDM-1_ in an *E. coli* isolate had partially identical sequences with a 39 kb IncR plasmid carrying *bla*_NDM-1_ and an 88 kb IncFII(pHN7A8)::IncN hybrid plasmid in a *C. freundii* isolate. 148–149 kb IncFIA(Hl1)::IncA/C2 plasmids and 75–76 kb IncFII(Yp) plasmids, both carrying *bla*_NDM-1_ were shared among three sequence type 11 (ST11) isolates and three ST395 isolates of *K. pneumoniae*, respectively. Most of the plasmids co-carried genes conferring resistance to clinically relevant antimicrobials, including third-generation cephalosporins, aminoglycosides, and fluoroquinolones, in addition to *bla*_NDM-1_. These results provide insight into the genetic basis of CRE in Vietnam, and could help control nosocomial infections.

## Introduction

Carbapenems have a broad spectrum of antimicrobial activity and are reserved for the treatment of infections caused by multidrug-resistant Gram-negative bacteria including Enterobacterales. The increase of cases infected by carbapenem-resistant Enterobacterales (CRE) is a serious threat to public health due to limited management of severe infections and risk of increased mortality [1,2]. Carbapenem-hydrolyzing β-lactamase (carbapenemase) genes, such as *bla*_NDM_, *bla*_KPC_, and *bla*_OXA-48_, that confer resistance to a broad range of β-lactams including third-generation cephalosporins and carbapenems are predominantly encoded on conjugative plasmids and have been transferred among Enterobacterales around the world [3]. There was a report that out of 27 carbapenem-resistant *Klebsiella pneumoniae* isolated from patients in a medical institution in Hanoi, Vietnam from 2014 to 2015, two isolates belonging to sequence type 395 (ST395) harbored *bla*_NDM-1_, and five isolates belonging to ST15, ST16, and ST2353 harbored *bla*_NDM-4_ [4]. There was another report that six *K. pneumoniae* isolates with *bla*_NDM-1_ were isolated from water samples in Hanoi, Vietnam in 2011 and belonging to ST283 [5]. Furthermore, we previously detected *bla*_NDM_ in 68.1% (47/69) of CRE isolates from 45 patients in a medical institution in Hanoi, Vietnam from 2010 to 2012, and some of isolates could be shown to transfer *bla*_NDM_ to *Escherichia coli* [6], suggesting that *bla*_NDM_ had spread via plasmids among Enterobacterales in medical institutions and communities in Vietnam.

The common replicon types of *bla*_NDM_-carrying plasmids associated with Enterobacterales includes IncA/C, IncFIA, IncFIB, IncFII, IncH, IncL/M, IncN, and IncX [7,8]. The plasmids occasionally co-carried other clinically relevant antimicrobial resistance (AMR) genes, such as extended-spectrum β-lactamase (ESBL) genes conferring resistance to third-generation cephalosporins, such as *bla*_CTX-M_, 16S ribosomal RNA (rRNA) methyltransferase genes conferring high-level resistance to aminoglycosides, such as *armA* and *rmt*, and fluoroquinolone resistance genes, such as *qnr* and *qep* [9]. Metallo-β -lactamase gene *bla*_NDM-1_ is often observed in a Tn*125* transposon in carbapenem-resistant *Acinetobacter* spp. Tn*125* is bracketed by two copies of insertion sequences, IS*Aba125*, belonging to the IS*30* family [10,11]. On plasmids of CRE, *bla*_NDM-1_ is also observed in the Tn*125*-like mobile gene element (MGE) which contains IS*Aba125* [11–14].

In this study, we report the detailed structures of *bla*_NDM-1_-carrying plasmids, and co-existing AMR genes and MGEs on the plasmids in Enterobacterales isolated in Vietnam. This knowledge provides insight into genetic characteristics and potential transmissions of the plasmids among Enterobacterales in Vietnam for the first time.

## Results

### Molecular characterization of carbapenem-nonsusceptible Enterobacterales isolated in Vietnam

A total of 122 isolates of ESBL-producing Enterobacterales were collected in daily diagnosis from individual patients in a medical institution in Hanoi, Vietnam between 2013 and 2017. Twenty-five Enterobacterales isolates, including *K. pneumoniae* (19/25, 76.0%), *E. coli* (2/25, 8.0%), *Citrobacter freundii* (2/25, 8.0%), *Morganella morganii* (1/25, 4.0%), and *Proteus mirabilis* (1/25, 4.0%) isolates, from clinical specimens, including blood (12/25, 48.0%), pus (10/25, 40.0%), urine (2/25, 8.0%), and sputum (1/25, 4.0%), were nonsusceptible to meropenem or imipenem and tested positive with CarbaNP test. The minimum inhibitory concentrations (MICs) of meropenem ranged from 1.5 to >32 μg/mL (median at 6 μg/mL) and those of imipenem ranged from 2 to >32 μg/mL (median at 12 μg/mL) (Fig. 1 and Table S1).

**Figure 1.**
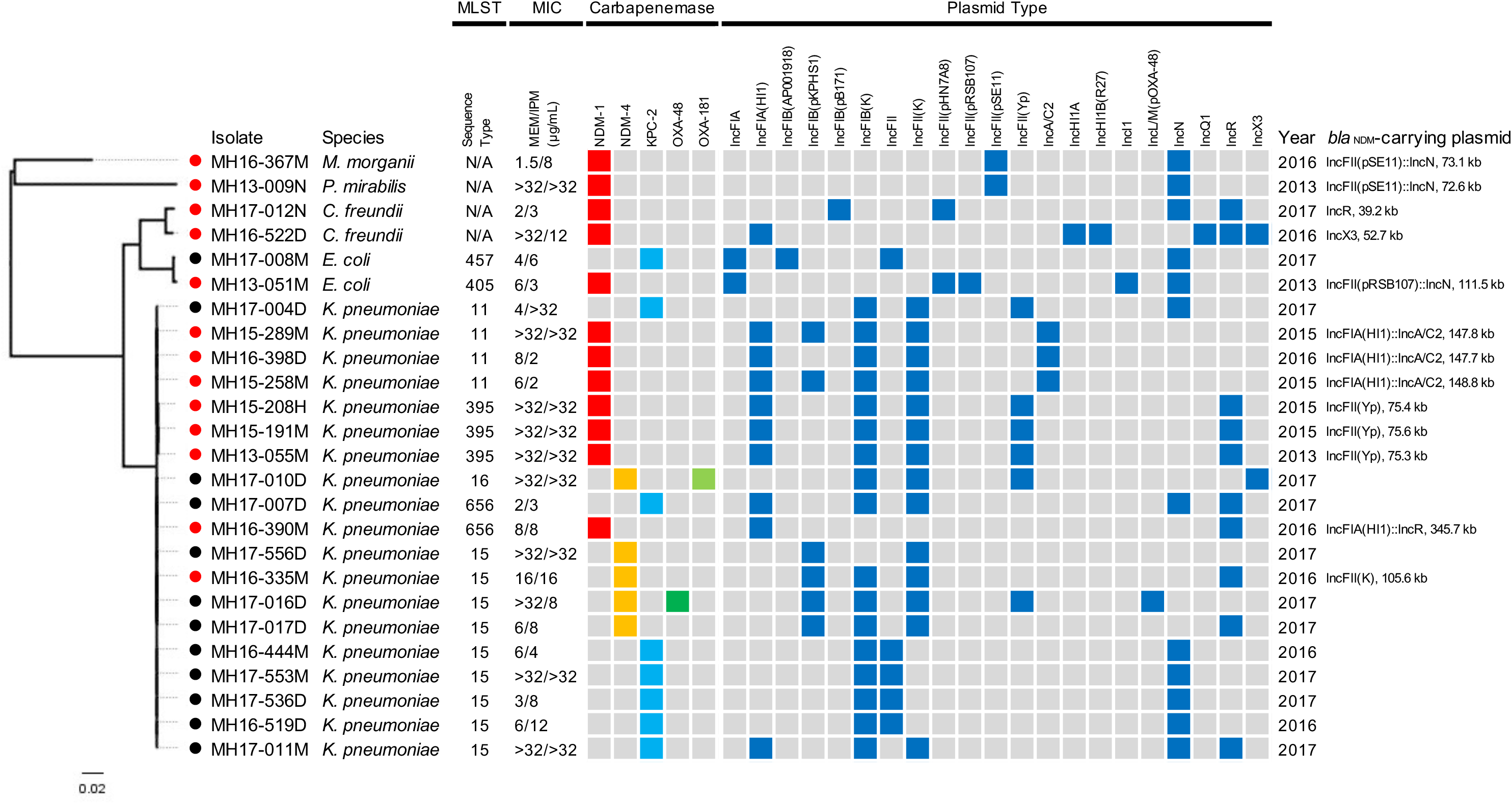
Carbapenem-nonsusceptible Enterobacterales isolates sequenced on Illumina systems. Red nodes, but not black nodes, show isolates subsequently sequenced with ONT MinION. Bar lengths represent the number of substitutions per site in the core genome constructed using the Roary pipeline (see Materials and Methods). Sequence types of multilocus sequence typing (MLST) analysis determined from genomes, minimum inhibitory concentrations (MICs) of meropenem (MEM) and imipenem (IPM) of isolates, carbapenemase genes detected by ResFinder and replicon types detected by PlasmidFinder in genomes, years in which the bacteria were isolated, and replicon types and sizes of *bla*_NDM_-carrying plasmids are shown.

Whole-genome sequencing of all 25 carbapenem-nonsusceptible Enterobacterales isolates with short-read sequencers resulted in the draft genome sequences ranged from 4.1 Mb to 5.9 Mb sizes with 132 to 868 contigs covered by at least x60 sequencing data (Table S1). Carbapenemase gene *bla*_NDM-1_ was detected in 12 Enterobacterales isolates (12/25, 48.0%), including *K. pneumoniae* (7/25, 28.0%), *C. freundii* (2/25, 8.0%), *E. coli* (1/25, 4.0%), *M. morganii* (1/25, 4.0%), and *P. mirabilis* (1/25, 4.0%) isolates. Carbapenemase gene *bla*_NDM-4_ was detected in five *K. pneumoniae* isolates (5/25, 20.0%), in which two isolates co-harbored other carbapenemase gene *bla*_OXA-48-like_ (*bla*_OXA-48_ and *bla*_OXA-181_, respectively) (2/25, 8.0%). Carbapenemase gene *bla*_KPC-2_ was detected in eight Enterobacterales isolates (8/25, 32.0%), including *K. pneumoniae* (7/25, 28.0%) and *E. coli* (1/25, 4.0%) isolates (Fig. 1 and Table S1).

Multilocus sequence typing (MLST) analysis revealed that the 19 carbapenemase-producing *K. pneumoniae* isolates belonged to five sequence types (STs), including ST15 (9/19, 47.4%), ST11 (4/19, 21.1%), ST395 (3/19, 15.8%), ST656 (2/19, 10.5%), and ST16 (1/19, 5.3%). The two *E. coli* isolates with *bla*_NDM-1_ and *bla*_KPC-2_ belonged to ST405 and ST457, respectively (Fig. 1). The presence of *bla*_NDM-1_ and replicons IncFII(pSE11) and IncN were detected in *M. morganii* MH16-367M and *P. mirabilis* MH13-009N, suggesting that these isolates might harbor *bla*_NDM-1_ on the identical IncFII(pSE11), IncN, or IncFII(pSE11)::IncN plasmid (Fig. 1).

MLST classified three *K. pneumoniae* isolates (MH15-289M, MH16-398D, and MH15-258M) as ST11 and another three *K. pneumoniae* isolates (MH15-208H, MH15-191M, and MH13-055M) as ST395. Interestingly, the presence of *bla*_NDM-1_ and replicons IncFIA(HI1), IncFIB(K), and IncFII(K) were detected in all six isolates (Fig. 1). In other examples, in the nine *K. pneumoniae*-ST15 isolates, *bla*_NDM-4_, IncFIB(pKPHS1), and IncFII(K) replicons were detected in four isolates (MH17-556D, MH16-335M, MH17-016D, and MH17-017D), and *bla*_KPC-2_, IncFIB(K), and IncN replicons were detected in five isolates (MH16-444M, MH17-553M, MH17-536D, MH16-519D, and MH17-011M) (Fig. 1).

### Putative transfer of *bla*_NDM-1_-carrying plasmids between Enterobacterales isolated in Vietnam

To investigate the similarities of the *bla*_NDM-1_-carrying plasmids in more detail, long-read sequencing was performed for 12 *bla*_NDM-1_-positive Enterobacterales isolates, including seven *K. pneumoniae* belonging to ST11 (three isolates), ST395 (three isolates), and ST656 (one isolate), two *C. freundii*, one *E. coli* belonging to ST405, one *M. morganii*, and one *P. mirabilis* isolates, and one *bla*_NDM-4_-positive *K. pneumoniae* isolate belonging to ST15 were further sequenced with the long-read sequencer. Hybrid analysis using short-read and long-read sequencing data resulted in complete sequences of 13 *bla*_NDM_-carrying plasmids that ranged from 39.2 kb to 345.7 kb sizes (covered by at least x125 long-read sequencing data; Table S2). Interestingly, many *bla*_NDM_-carrying plasmids co-carried other clinically relevant AMR genes, such as 16S rRNA methyltransferases genes (9/13, 69.2%) and quinolone resistance genes (6/13, 46.2%) on the same plasmid (Table S2).

BLAST search revealed that several *bla*_NDM-1_-carrying plasmids had high sequence identities with each other and with plasmids in the GenBank database (Figs. 2, 3, 4, and 5). However, *bla*_NDM-1_-carrying plasmid pMH16-522D_1 (52.7 kb plasmid IncX3 replicon, accession: AP018571) in *C. freundii* MH16-522D, *bla*_NDM-1_-carrying plasmid pMH16-390M_1 [345.7 kb plasmid with IncFIA(HI1)::IncR replicons, accession: AP018583] in *K. pneumoniae* MH16-390M, and *bla*_NDM-4_-carrying plasmid pMH16-335M_1 [105.6 kb plasmid with IncFII(K) replicon, accession: AP018584] in *K. pneumoniae* MH16-335M were not homologous to any other plasmids in this study (Table S2). BLAST search revealed that numerous plasmids in Enterobacterales isolated around the world had 98-100% identity in more than 70% of the regions of pMH16-522D_1, pMH16-390M_1, and pMH16-335M_1, respectively. All three plasmids harbored additional β -lactamase and/or *qnr* genes: *C. freundii* pMH16-522D_1 carried *bla*_SHV-12_, *K. pneumoniae* pMH16-390M_1 carried *bla*_CTX-M-3_, *bla*_TEM-1b_, and *qnrB6*, and *K. pneumoniae* pMH16-335M_1 carried *bla*_OXA-9_, *bla*_SHV-106_, and *bla*_SHV-28_ (Table S2). Furthermore, all three plasmids contained the Tn*125*-like MGEs around *bla*_NDM_, and the sequences were highly identical to the reference sequence of *Tn*125 (accession: HQ857107) (96.9 to 100% identity with 87-89% region) [14].

**Figure 2.**
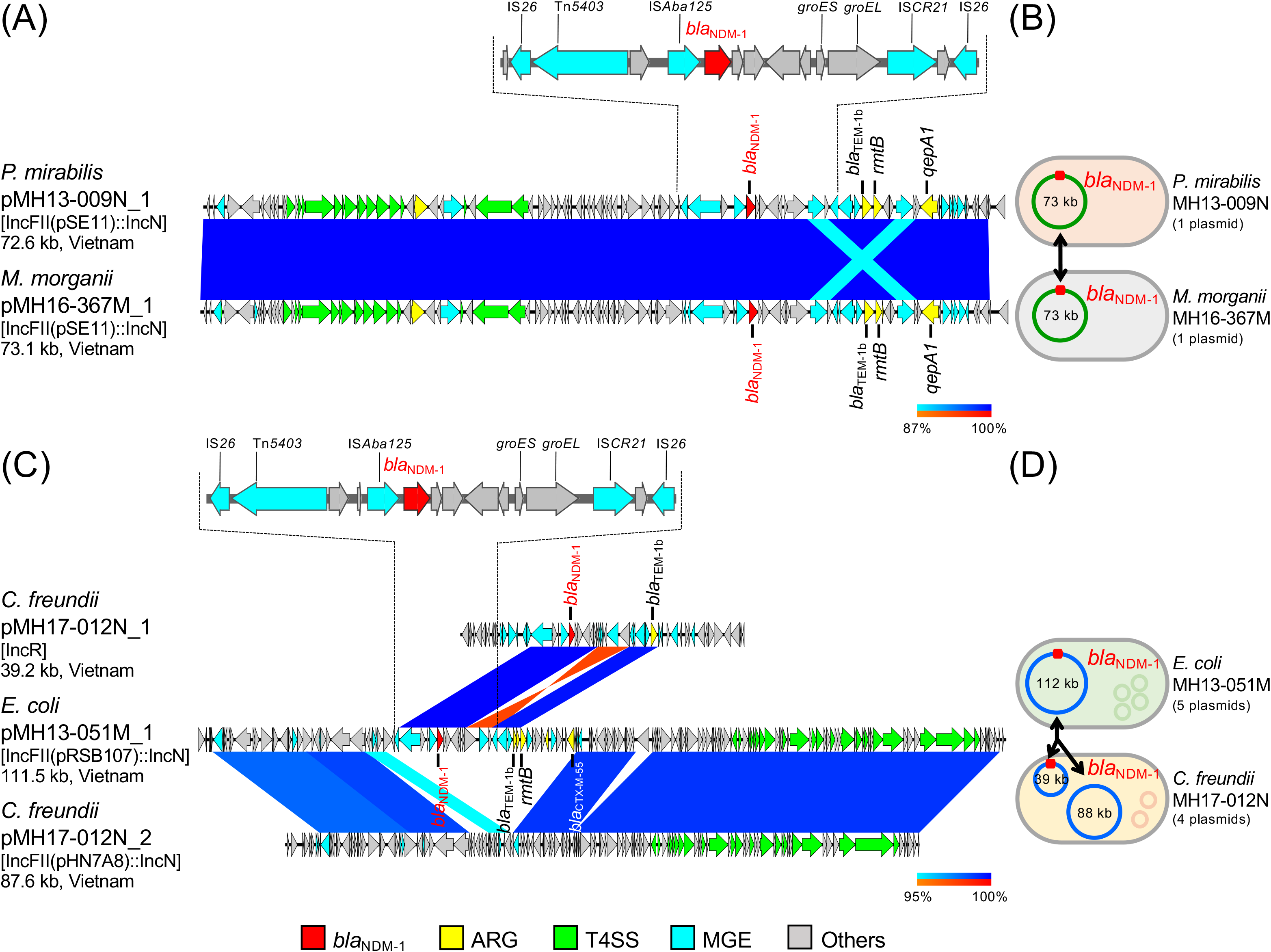
*bla*_NDM-1_-carrying plasmids and the detailed genetic structures around *bla*_NDM-1_ in carbapenem-nonsusceptible Enterobacterales isolates sequenced using ONT MinION and Illumina systems. Sets of plasmids from *P. mirabilis* MH13-009N (pMH13-009N_1, accession: AP018566) and *M. morganii* MH16-367M (pMH16-367M_1, accession: AP018565), and from *E. coli* MH13-051M (pMH13-051M_1, accession: AP018572) and *C. freundii* MH17-012N (pMH17-012N_1, accession: AP018567 and pMH17-012N_2, accession: AP018568), are shown. (A) and (C) Linear comparison of *bla*_NDM-1_-carrying plasmid sequences. (B) and (D) Schematic representation of possible transfer of plasmids, including *bla*_NDM-1_-carrying plasmids. Circles within bacterial cells show plasmids, and red squares on plasmids show *bla*_NDM-1_. Sets of plasmids from *P. mirabilis* MH13-009N and *M. morganii* MH16-367M (A and B), and from *E. coli* MH13-051M and *C. freundii* MH17-012N (C and D), are shown. Red, yellow, green, blue, and gray arrows indicate *bla*_NDM_, other important AMR genes (ARGs), type IV secretion system (T4SS)-associated genes involved in conjugation, mobile gene elements (MGEs), and other genes, respectively. The colors in comparison of plasmids show percent identity and sequence direction as indicated. Blue for matches in the same direction and red for matches in the inverted direction.

**Figure 3.**
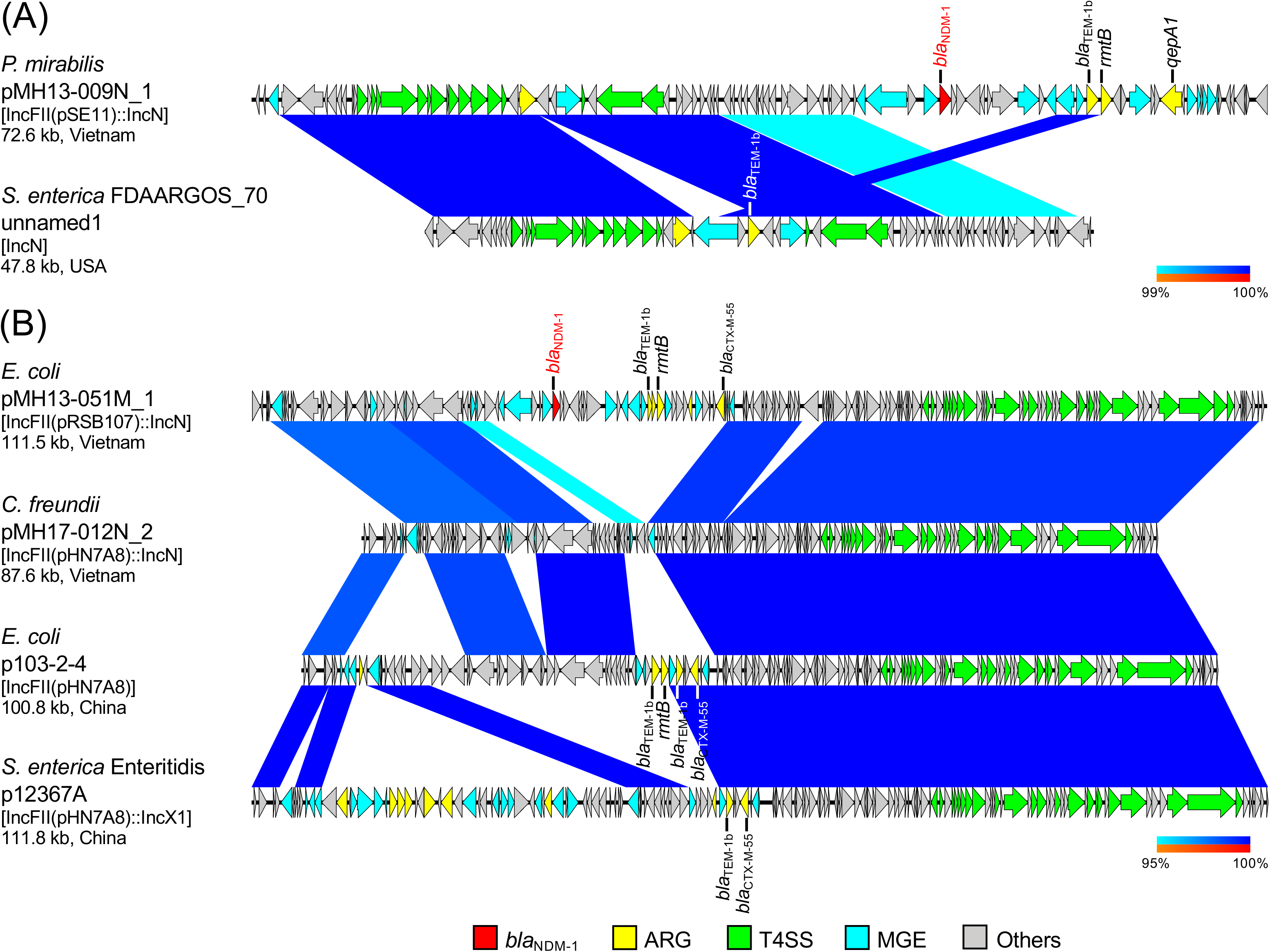
(A) and (B) Linear comparison of plasmid sequences from Vietnam and other countries. *P. mirabilis* pMH13-009N_1 (this study, accession: AP018566) and *Salmonella enterica* FDAARGOS_70 plasmid unnamed1 (accession: CP026053) (A), and *E. coli* pMH13-051M_1 (this study, accession: AP018572), *C. freundii* pMH17-012N_2 (this study, accession: AP018568), *E. coli* p103-2-4 (accession: CP034846), and *Salmonella enterica* serovar Enteritidis p12367A (accession: CP041177) (B) are shown. Red, yellow, green, blue, and gray arrows indicate *bla*_NDM_, other important AMR genes (ARGs), type IV secretion system (T4SS)-associated genes involved in conjugation, mobile gene elements (MGEs), and other genes, respectively. The colors in comparison of plasmids show percent identity and sequence direction as indicated. Blue for matches in the same direction and red for matches in the inverted direction.

**Figure 4.**
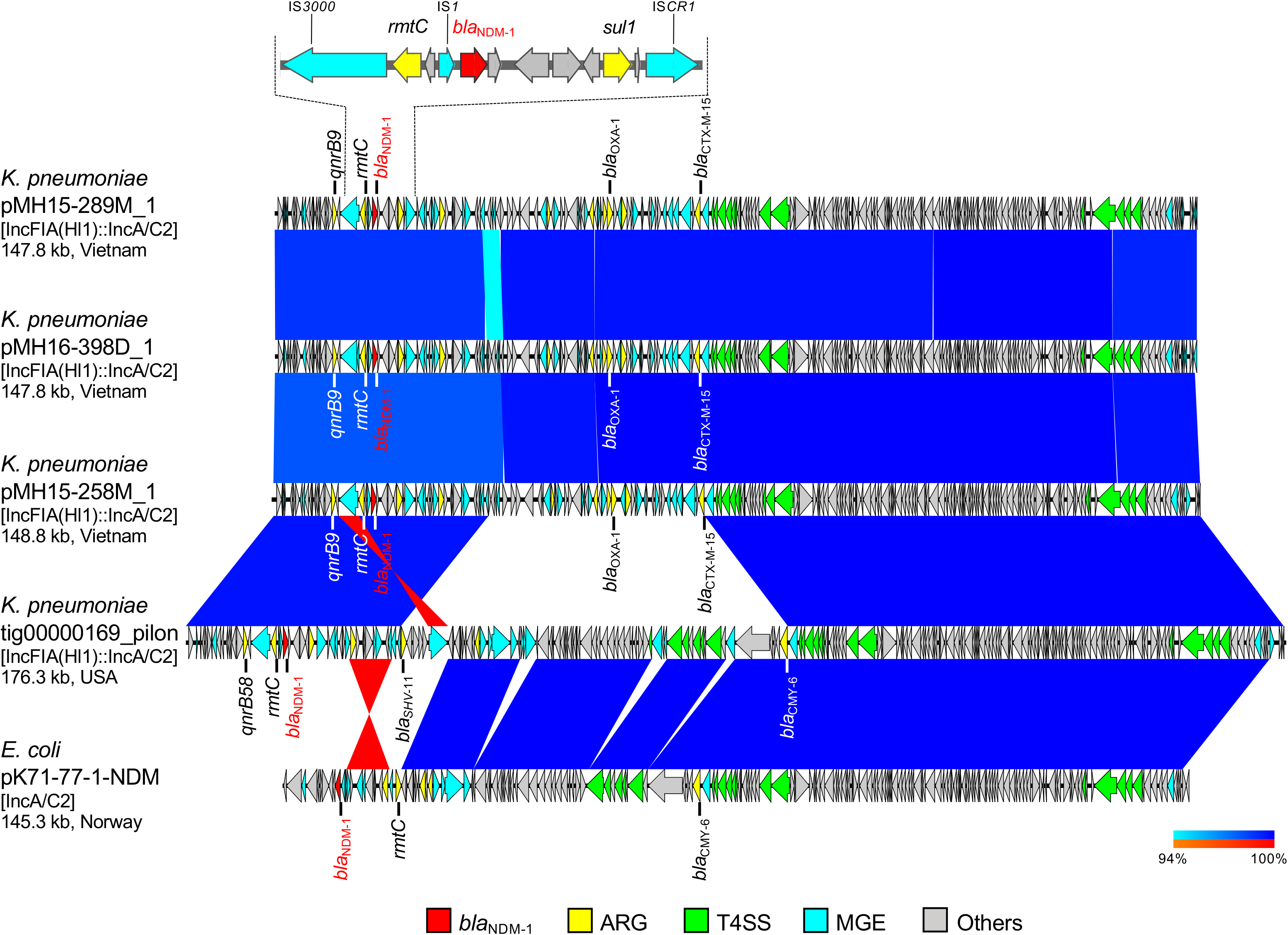
Linear comparison of *bla*_NDM-1_-carrying plasmid sequences and the detailed genetic structures around *bla*_NDM-1_ from Vietnam and other countries. *K. pneumoniae* pMH15-289M_1 (this study, accession: AP018577), *K. pneumoniae* pMH16-398D_1 (this study, accession: AP018578), *K. pneumoniae* pMH15-258M_1 (this study, accession: AP018579), *K. pneumoniae* tig00000169_pilon (accession: CP021952), and *E. coli* pK71-77-1-NDM (accession: CP040884) are shown. Red, yellow, green, blue, and gray arrows indicate *bla*_NDM_, other important AMR genes (ARGs), type IV secretion system (T4SS)-associated genes involved in conjugation, mobile gene elements (MGEs), and other genes, respectively. The colors in comparison of plasmids show percent identity and sequence direction as indicated. Blue for matches in the same direction and red for matches in the inverted direction.

**Figure 5.**
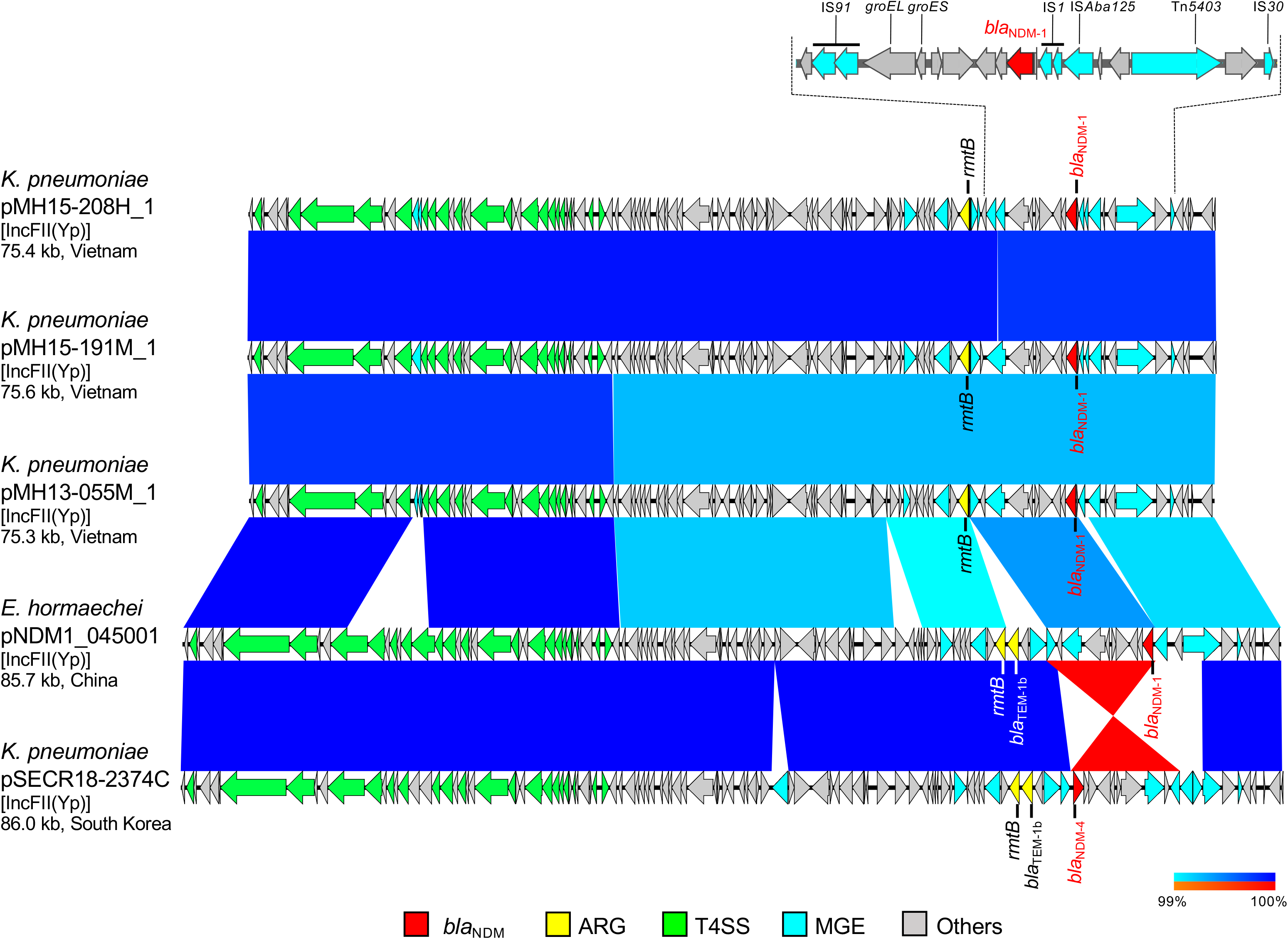
Linear comparison of *bla*_NDM_-carrying plasmid sequences and the detailed genetic structures around *bla*_NDM-1_ from Vietnam and other countries. *K. pneumoniae* pMH15-208H_1 (this study, accession: AP018580), *K. pneumoniae* pMH15-191M_1 (this study, accession: AP018581), *K. pneumoniae* pMH13-055M_1 (this study, accession: AP018582), *E. hormaechei* pNDM1_045001 (accession: CP043383), and *K. pneumoniae* pSECR18-2374C (accession: CP041930) are shown. Red, yellow, green, blue, and gray arrows indicate *bla*_NDM_, other important AMR genes (ARGs), type IV secretion system (T4SS)-associated genes involved in conjugation, mobile gene elements (MGEs), and other genes, respectively. The colors in comparison of plasmids show percent identity and sequence direction as indicated. Blue for matches in the same direction and red for matches in the inverted direction.

*P. mirabilis* MH13-009N (isolated in 2013) and *M. morganii* MH16-367M (isolated in 2016) had only one plasmid, pMH13-009N_1 [72.6 kb plasmid with IncFII(pSE11)::IncN replicons, accession: AP018566] and pMH16-367M_1 [73.1 kb plasmid with IncFII(pSE11)::IncN replicons, accession: AP018565], respectively; the sequences of these plasmids were almost identical (100% identity in 99% regions), including the *bla*_NDM-1_ and IncFII(pSE11)::IncN hybrid replicon regions (Figs. 2A and 2B). Both of the plasmids carried *bla*_NDM-1_ and an integron cassette that contains the class 1 integrase gene (*intI1*) and the AMR genes, such as *bla*_TEM-1b_, *rmtB*, and *qepA1*. The same integron cassette was observed on *E. coli* pHPA (10.5 kb plasmid with IncFII replicon, accession: AB263754) isolated from a human in Japan in 2002 [15]. Carbapenemase gene *bla*_NDM-1_ was surrounded by several MGEs, including IS*Aba125*, IS*CR21*, and two IS*26*, on *P. mirabilis* pMH13-009N_1 and *M. morganii* pMH16-367M_1, and shared identical sequences with Tn*125* (Fig. 2A): 100% identity with 89% region of the reference sequence (accession: HQ857107) [14,16,17]. The nearly identical *bla*_NDM-1_-carrying plasmids, *P. mirabilis* pMH13-009N_1 and *M. morganii* pMH16-367M_1 were successfully transferred to recipient *E. coli* at a frequency of 10^−2^–10^−3^ *in vitro*.

*E. coli* MH13-051M (isolated in 2013) harbored *bla*_NDM-1_ on plasmid pMH13-051M_1 [111.5 kb plasmid with IncFII(pRSB107)::IncN replicons, accession: AP018572], which had partially identical sequences with two plasmids, pMH17-012N_1 (39.2 kb plasmid with IncR replicon, accession: AP018567, 100% identity in 24% region of pMH13-051M_1) and pMH17-012N_2 [87.6 kb plasmid with IncFII(pHN7A8)::IncN replicons, accession: AP018568, 98.8% identity in 78% region of pMH13-051M_1], both found in *C. freundii* MH17-012N (isolated in 2017) (Figs. 2C and 2D). *E. coli* pMH13-051M_1 [IncFII(pRSB107)::IncN plasmid] and *C. freundii* pMH17-012N_1 (smaller IncR plasmid) commonly carried *bla*_NDM-1_ and *bla*_TEM-1b_, and pMH13-051M_1 further carried *bla*_CTX-M-55_ and *rmtB*. *C. freundii* pMH17-012N_2 [larger IncFII(pHN7A8)::IncN plasmid] carried no known AMR genes, but did have a set of conjugation-associated type IV secretion system (T4SS) genes, which shared identical sequences with *E. coli* pMH13-051M_1. The genetic structures surrounding *bla*_NDM-1_ in *E. coli* pMH13-051M_1 and *C. freundii* pMH17-012N_1 contained MGEs, including IS*Aba125*, IS*CR21*, and two IS*26*, and shared identical sequences with Tn*125* (Fig. 2C): 98.6% and 100% identity with 89% region of the reference sequence (accession: HQ857107) [14].

### Comparison of *bla*_NDM-1_-carrying plasmids in Vietnam with those in other countries

Next, we compared *bla*_NDM-1_–carrying plasmids identified in Vietnam with those reported in other countries. A set of conjugation-associated T4SS genes in *P. mirabilis* pMH13-009N_1 [72.6 kb plasmid with IncFII(pSE11)::IncN replicons] was partially identical with *Salmonella enterica* FDAARGOS_70 plasmid unnamed1 (47.8 kb plasmid with IncN replicon, accession: CP026053, 100% identity in 61% region of pMH13-009N_1) isolated from a human in the United States in 2013, which carried *bla*_TEM-1b_ (Fig. 3A). However, *S. enterica* FDAARGOS_70 plasmid unnamed1 did not carry *bla*_NDM-1_ as *P. mirabilis* pMH13-009N_1 did. *C. freundii* pMH17-012N_2 [87.6 kb plasmid with IncFII(pHN7A8)::IncN replicons and no AMR genes] was highly identical with *E. coli* p103-2-4 [100.8 kb plasmid with IncFII(pHN7A8) replicons, accession: CP034846, 99.9% identity in 94% region of pMH17-012N_2] from a goose farm in China in 2018 and with *S. enterica* serovar Enteritidis p12367A [111.8 kb plasmid with IncFII(pHN7A8)::IncX1 replicons, accession: CP041177, 99.9% identity in 73% region of pMH17-012N_2] isolated from a human in China in 2013 (Fig. 3B). *E. coli* p103-2-4 and *S. enterica* serovar Enteritidis p12367A carried several AMR genes, such as *bla*_CTX-M-55_ and *bla*_TEM-1b_.

Furthermore, three *K. pneumoniae* isolates belonging to ST11 (MH15-289M, MH16-398D, and MH15-258M isolated in 2015, 2016, and 2015, respectively) shared nearly identical *bla*_NDM-1_-carrying plasmids, pMH15-289M_1, pMH16-398D_1, and pMH15-258M_1 [147–149 kb plasmids with IncFIA(Hl1)::IncA/C2 replicons, accessions: AP018577, AP018578, and AP018579, respectively, 99.6% identity in 99% regions], and all of these plasmids carried other AMR genes, such as *bla*_CTX-M-15_, *bla*_OXA-1_, *rmtC*, and *qnrB9* (Fig. 4). *bla*_NDM-1_ was surrounded by several MGEs, including IS*3000*, IS*1*, and IS*CR1*. These three plasmids were highly identical with *K. pneumoniae* plasmid tig00000169_pilon [176.3 kb plasmid with IncFIA(Hl1)::IncA/C2 replicons, accession: CP021952, 99.8% identity in 82% region of pMH15-289M_1] in the United States and with *E. coli* pK71-77-1-NDM (145.3 kb plasmid with IncA/C2 replicon, accession: CP040884, 99.9% identity in 62% region of pMH15-289M_1) isolated from a human in Norway in 2010. The plasmids tig00000169_pilon and pK71-77-1-NDM had identical sequences in the T4SS region of pMH15-289M_1, and carried *bla*_NDM-1_, *bla*_CMY-6_, and *rmtC*, and the plasmid tig00000169_pilon further carried *bla*_SHV-11_ and *qnrB58*. *K. pneumoniae* MH16-398D had 235 sequence variants (214 single-nucleotide variants [SNVs], 14 multiple-nucleotide variants [MNVs], four deletions, two insertions, and one replacement), and *K. pneumoniae* MH15-258M had six sequence variants (five SNVs and one replacement) relative to *K. pneumoniae* MH15-289M.

Another three *K. pneumoniae* isolates belonging to ST395 (MH15-208H, MH15-191M, and MH13-055M isolated in 2015, 2015, and 2013, respectively) shared nearly identical *bla*_NDM-1_-carrying plasmids, pMH15-208H_1, pMH15-191M_1, and pMH13-055M_1 [75–76 kb plasmids with IncFII(Yp) replicon, accessions: AP018580, AP018581, and AP018582, respectively, 99.8–99.9% identity in 99% regions], and all plasmids carried another AMR gene, *rmtB* (Fig. 5). *bla*_NDM-1_ in these plasmids was surrounded by several MGEs, including IS*Aba125*, IS*1*, and IS*91*, and shared partially identical sequences with Tn*125* (Fig. 5): 99.5-100% identity with 88-89% region of the reference sequence (accession: HQ857107) [14]. These three plasmids were highly identical with *Enterobacter hormaechei* pNDM1_045001 [85.7 kb plasmid with IncFII(Yp) replicon, accession: CP043383, 99.9% identity in 98% region of pMH15-208H_1] isolated from a human in China in 2018 and with *K. pneumoniae* pSECR18-2374C [86.0 kb plasmid with IncFII(Yp) replicon, accession: CP041930, 99.9% identity in 95% region of pMH15-208H_1] isolated from a human in South Korea in 2018. *E. hormaechei* pNDM1_045001 and *K. pneumoniae* pSECR18-2374C had a partially identical sequence of conjugation-associated T4SS genes with pMH15-208H_1. *E. hormaechei* pNDM1_045001 carried *bla*_NDM-1_, *bla*_TEM-1b_, and *rmtB*, and *K. pneumoniae* pSECR18-2374C carried *bla*_NDM-4_, *bla*_TEM-1b_ and *rmtB*. *K. pneumoniae* MH15-191M had three sequence variants (two SNVs and one MNV) and *K. pneumoniae* MH13-055M had only one SNV relative to *K. pneumoniae* MH15-208H.

In bacterial conjugation experiments with *E. coli*, *C. freundii* pMH17-012N_1 (39.2 kb plasmid with IncR replicon), pMH17-012N_2 [87.6 kb plasmid with IncFII(pHN7A8)::IncN replicons], *E. coli* pMH13-051M_1 [111.5 kb plasmid with IncFII(pRSB107)::IncN replicons], *K. pneumoniae* pMH15-258M_1 [148.8 kb plasmid with IncFIA(Hl1)::IncA/C2 replicons], and *K. pneumoniae* pMH15-208H_1 [75.4 kb plasmid with IncFII(Yp) replicon] were not transferred to recipient *E. coli* under our experimental conditions though a set of T4SS-associated genes were detected in *C. freundii* pMH17-012N_2, *E. coli* pMH13-051M_1, *K. pneumoniae* pMH15-258M_1, and *K. pneumoniae* pMH15-208H_1.

## Discussion

The Viet Nam Resistance (VINARES), a nationwide surveillance network consisting of 16 central and provincial-level medical institutions was established in Vietnam in 2013 [18]. According to VINARES data, the rates of resistance of *K. pneumoniae* to third-generation cephalosporins, carbapenems, aminoglycosides, and fluoroquinolones were 66.4%, 17.1%, 29.5%, and 53.0%, respectively, in Vietnam [19], whereas the rates were 13.6%, 0.5%, 6.5%, and 8.7%, respectively, in the United Kingdom in 2013 [20], and 17.2%, 4.3%, 4.5%, and 16.8%, respectively, in the United States in 2010 [21]. Though the rates of resistance of Enterobacterales to major antimicrobials are much higher in Vietnam than in Western countries, the genetic causes remain unknown.

In this study, we performed comprehensive genetic analysis on plasmids carrying *bla*_NDM_ carbapenemase genes in Enterobacterales isolated from patients in Vietnam. We focused on carbapenem-nonsusceptible Enterobacterales isolates harboring *bla*_NDM_, one of the most prevalent carbapenemase genes in the world, and completely sequenced 12 plasmids carrying *bla*_NDM-1_ from *K. pneumoniae* (ST11, ST395, and ST656), *E. coli* (ST405), *C. freundii*, *M. morganii*, and *P. mirabilis* isolates and one plasmid carrying *bla*_NDM-4_ from *K. pneumoniae* isolate (ST15). Nearly identical plasmids carrying *bla*_NDM-1_ were detected in different bacterial species of this study, suggesting that they represent common and important AMR plasmids disseminated in Vietnam (Figs. 2, 4, and 5). Moreover, the co-existence of clinically relevant AMR genes, such as ESBL genes (e.g., *bla*_CTX-M_, *bla*_OXA_, and *bla*_TEM_), aminoglycoside resistance genes (e.g., *armA* and *rmt*), and fluoroquinolone resistance genes (e.g., *qep* and *qnr*), with *bla*_NDM-1_ was observed in several plasmids, such as *P. mirabilis* pMH13-009N_1 (Fig. 2) and *K. pneumoniae* pMH15-289M_1 (Fig. 4). Hence, the spread of the plasmids both within and among species of bacteria would have caused an increase in multidrug-resistant bacteria and the infected hosts in Vietnam, and will pose a threat to public health.

Almost identical IncFII(pSE11)::IncN hybrid plasmids, pMH13-009N_1 and pMH16-367M_1, were found in *P. mirabilis* and *M. morganii* isolates, respectively (Figs. 2A and 2B). These plasmids had identical sequences including a set of conjugation-associated T4SS genes with the IncN plasmid from *S. enterica* isolated in the United States (Fig. 3A). IncN plasmids are prevalent in the microbiota of animals, and disseminate AMR genes such as *bla*_CTX-M-1_ and *qnr* [22], suggesting that IncN plasmids with the same origin of *P. mirabilis* pMH13-009N_1 and *M. morganii* pMH16-367M_1 could be disseminated in communities, including humans, animals, and the environment, in Vietnam and other countries.

The IncFII(pRSB107)::IncN hybrid plasmid pMH13-051M_1 in *E. coli*, had partial sequence identity with the IncR plasmid pMH17-012N_1 and the IncFII(pHN7A8)::IncN hybrid plasmid pMH17-012N_2 both in *C. freundii* (Figs. 2C and 2D). Rearrangement of conjugation genes plays an important role in the dissemination of AMR genes between bacteria [23]. Because known IncR plasmids are non-transferable [24], an IncR plasmid carrying AMR genes (e.g. *C. freundii* pMH17-012N_1) and another plasmid carrying conjugation genes (e.g. *C. freundii* pMH17-012N_2) could be assumed to have been fused into a single plasmid with both AMR and conjugation genes (e.g. *E. coli* pMH13-051M_1) in a bacterium. Subsequently the plasmid was propagated by conjugation to another bacterium. There is an increasing number of reports of IncR plasmids carrying various AMR genes and the pool of AMR genes on IncR plasmids is thought to spread to transmissible plasmids via recombination, contributing to the high evolutionary plasticity of bacterial genomes [25]. Further analysis on the possible recombination events are needed to understand the ways of transmission of these plasmids between bacterial isolates and species.

The IncFII(pHN7A8)::IncN hybrid plasmid pMH17-012N_2 had highly identical sequences including a set of conjugation-associated T4SS genes with the plasmids *E. coli* p103-2-4 and *S. enterica* Enteritidis p12367A, that carry several AMR genes, such as *bla*_CTX-M-55_ and *bla*_TEM-1b_, but no *bla*_NDM_ (Fig. 3B). The IncFII plasmid pHN7A8 (accession: JN232517), which also carries several AMR genes, such as *bla*_CTX-M-65_, *rmtB*, and *fosA3*, and is widespread in *E. coli* from farm and companion animals in China [26], suggesting that plasmids with IncFII(pHN7A8) replicon are associated with AMR.

*bla*_NDM-1_ was found on several hybrid plasmids, such as IncFII(pSE11)::IncN, IncFII(pRSB107)::IncN hybrid plasmids (Figs. 2A and 2C), and IncFIA(Hl1)::IncA/C2 hybrid plasmids from three *K. pneumoniae* ST11 isolates (Fig. 4). Hybrids of multiple replicons belonging to different incompatibility groups represent a plasmid strategy for expansion of host range and dissemination of acquired AMR genes among bacteria [27]. The IncFIA(Hl1)::IncA/C2 hybrid plasmids or IncFII(Yp) plasmids were reported from other regions, including the United States, Europe, and Asia (Figs. 4 and 5). IncFIA(Hl1)::IncA/C2 plasmids and IncFII(Yp) plasmids were detected in *K. pneumoniae* in this study, suggesting that these plasmids and/or bacterial isolates would have also been spreading in medical institutions and community in Vietnam.

The *bla*_NDM-1_-surrounding regions in plasmids, *P. mirabilis* pMH13-009N_1, *M. morganii* pMH16-367M_1, *C. freundii* pMH17-012N_1, and *E. coli* pMH13-051M_1 (Fig. 2), *K. pneumoniae* pMH15-208H_1, pMH15-191M_1, and pMH13-055M_1 (Fig. 5), and *K. pneumoniae* pMH16-390M_1, pMH16-335M_1, and *C. freundii* pMH16-522D_1 had almost identical sequence including Tn*125* which was discovered in *A. baumannii* [10,14,16]. Tn*125* includes IS*Aba125* and IS*CR21*, but sequences of the Tn*125*-like region in *P. mirabilis* pMH13-009N_1, *M. morganii* pMH16-367M_1, and *E. coli* pMH13-051M_1 (Fig. 2) include two IS*26* in addition to IS*Aba125* and IS*CR21*. This Tn*125*-like structure has also been detected in other plasmids from Enterobacterales, such as *C. freundii* pCF104a-T3 (accession: MN657241) isolated from a human in Germany in 2015 [17]. This report showed that IS*26*-mediated transfer of *bla*_NDM-1_-containing Tn*125*-like regions had occurred in different Enterobacterales species within one medical institution. The MGEs surrounding *bla*_NDM-1_ on plasmids, *K. pneumoniae* pMH15-289M_1, pMH16-398D_1, and pMH15-258M_1 (Fig. 4), were highly identical with the corresponding sequence of *E. coli* pEC2-NDM-3 (accession: KC999035) isolated from a human in Australia in 2010 though pEC2-NDM-3 carried *bla*_NDM-3_ with a part of the same set of MGEs, including IS*CR1*. IS*CR1* was suggested to provide a vehicle for *bla*_NDM-1_ dissemination [8, 28].

MLST analysis is helpful to estimate whether bacterial isolates belonging to the same STs were originated from the same clones and spread within the medical institution, and whole-genome SNV-based analysis enables us to estimate more accurate genetic relationship. According to a report [29], SNVs accumulated in *K. pneumoniae* ST258 isolates at a rate of 3.9 SNVs per year. Our analysis of SNVs of *K. pneumoniae* ST11 isolates (Fig. 4) revealed 214 SNVs in MH16-398D relative to MH15-289M. This suggests that different ST11 clones are present in the medical institutions and communities. In support of this theory, a report [30] showed that *K. pneumoniae* isolates from medical institutions, sewage, canals, and agricultural waste water were intermixed in the phylogenetic classification. Because ST11 is one of major international epidemic clones of *K. pneumoniae* [31], an ST11 clone carrying AMR plasmid could have disseminated in a medical institution or local community in Hanoi, Vietnam. Another three ST395 isolates of *K. pneumoniae* (Fig. 5) are very clonal based on the rate of SNV accumulation [29] and could be disseminated from the same clone within the medical institution because only a few SNVs were present in MH15-191M and MH13-055M relative to MH15-208H.

## Conclusion

Plasmid analysis in this study shows the complex structures and diversity of plasmids carrying *bla*_NDM-1_ in Vietnam. Hybrid analysis with both Illumina short-read and ONT long-read sequencing is a promising method for detecting important AMR plasmids in CRE isolates and the real-time analysis is useful for controlling nosocomial infections.

## Materials and Methods

### Bacterial isolates

A total of 122 ESBL-producing Enterobacterales isolates were collected from selected specimens, including blood, sputum, urine, and pus from both in-patients and out-patients with infections in daily diagnosis in a reference medical institution in Hanoi, Vietnam between 2013 and 2017. All bacterial isolates were excluded from duplicates from the same patient. The medical institution has 700 beds, and includes 13 surgical departments and 10 internal medicine departments. The medical services are provided for both military and civil patients. Bacterial species identification, prediction of ESBL activity, and antimicrobial susceptibility testing were performed by Vitek 2 (BioMèrieux) and E-test (BioMèrieux). ESBL gene(s)-harboring isolates could also harbor other important AMR genes, including carbapenemase gene(s). According to the Clinical and Laboratory Standards Institute (CLSI) 2020 guidelines, breakpoints of meropenem and imipenem are ≤1 μg/mL (susceptible), 2 μg/mL (intermediate), ≥4 μg/mL (resistant), respectively. Carbapenemase activity was examined by CarbaNP test according to the CLSI 2020 guidelines. Major carbapenemase genes (*bla*_NDM_, *bla*_KPC_, *bla*_OXA-48_, *bla*_IMP_, and *bla*_VIM_) were detected by a multiplex PCR method [32]. Twenty-five carbapenem-nonsusceptible and carbapenemase-positive isolates (19 *K. pneumoniae*, two *E. coli*, two *C. freundii*, one *M. morganii*, and one *P. mirabilis*) were subjected to the whole-genome sequencing analysis described below.

### Preparation of genomic DNA

For short-read sequencing, genomic DNAs of bacterial isolates were extracted with the phenol-chloroform method, and purified with QIAquick PCR Purification Kit (QIAGEN) according to the manufacturer’s instructions. For long-read sequencing, high-molecular-weight genomic DNAs of bacterial isolates were extracted with MagAttract HMW DNA Kit (QIAGEN) according to the manufacturer’s instructions. The extracted genomic DNAs were quantified with a Qubit 2.0 fluorometer (Thermo Fisher Scientific).

### Whole-genome sequencing and bioinformatics analysis

Whole-genome sequencing was performed to examine the prevalence of important AMR genes on plasmids. Long-read sequencing is useful for constructing complete sequences of whole plasmids and tracking horizontal transfer of AMR plasmids during nosocomial infections [33]. Recently, long-read nanopore sequencing using MinION nanopore sequencer from Oxford Nanopore Technologies (ONT) has been applied to such investigations [34,35].

Whole genome sequencing using MiniSeq or HiSeq systems (Illumina) was performed for phylogenetic and MLST analysis, and screening of acquired AMR genes and plasmid replicon types. Library for Illumina sequencing (insert size of 500-900 bp) was prepared using Nextera XT DNA Library Prep Kit (Illumina). Paired-end sequencing was performed using MiniSeq (2 × 150 bp) or HiSeq 4000 systems (2 × 150 bp). Trimming and *de novo* assembly of paired-end reads was performed using CLC Genomics Workbench v12.0 (QIAGEN) with default parameters. A maximum-likelihood phylogenetic tree was generated by PhyML from a concatenated core-gene alignment consisting of 13,305 SNVs constructed using the Roary pipeline (https://sanger-pathogens.github.io/Roary/). For PhyML, we used the following parameters that indicate the GTR +G4 model of DNA substitution with estimation of the shape parameter of the gamma distribution by maximizing the likelihood: -m GTR -c 4 -a e.

Twelve Enterobacterales isolates with *bla*_NDM-1_ (seven *K. pneumoniae*, one *E. coli*, two *C. freundii*, one *M. morganii*, and one *P. mirabilis*) and one *K. pneumoniae* isolate with *bla*_NDM-4_ were further sequenced on MinION (ONT) using the SQK-RAD002 or SQK-RBK001 kits and R9.4 flowcells according to the manufacturer’s instructions to obtain complete sequences of plasmids carrying carbapenem resistance genes. *De novo* assembly was performed using Canu v1.5 [36] and Miniasm [37] with default parameters. The overlap region in the assembled contig was detected by genome-scale sequence comparison using LAST (http://last.cbrc.jp) and was trimmed manually. Illumina paired-end reads were mapped onto the resulted circular sequences, and error correction was performed by extracting consensus of mapped reads using CLC Genomics Workbench v12.0 with default parameters.

Sequence types, carbapenemase genes, and plasmid replicon types were analyzed using MLST v2.0, ResFinder v3.2 with minimum threshold of 90% identity and 60% coverage, and PlasmidFinder v2.1 with minimum threshold of 90% identity and 60% coverage, respectively, on the Center for Genomic Epidemiology (CGE) server at Technical University of Denmark (http://www.genomicepidemiology.org).

Coding sequences (CDS) annotation using RASTtk on the PATRIC v3.6.3 server (https://www.patricbrc.org) with default parameters. Sequence variant detection was performed using CLC Genomics Workbench v12.0 with default parameters of Fixed Ploidy Variant Detection (Ploidy=1). Linear comparison of sequences was performed using BLAST with the default settings (the nucleotide collection database and the megablast program), and visualized by Easyfig (http://mjsull.github.io/Easyfig/). *bla*_NDM-1_, other important AMR genes (ARG), type IV secretion system (T4SS)-associated genes involved in conjugation that were detected by SecReT4 program [38], and mobile gene elements (MGEs) were identified manually from CDS annotations and basically analyzed by comparing the sequences analyzed in previous studies.

Draft genome and complete plasmid sequences of carbapenem-nonsusceptible Enterobacterales isolated in Vietnam have been deposited at GenBank/EMBL/DDBJ under BioProject number PRJDB6655.

### Bacterial conjugation

Bacterial conjugation was performed using six *bla*_NDM-1_-carrying plasmid-positive Enterobacterales isolates [*P. mirabilis* MH13-009N, *M. morganii* MH16-367M, *C. freundii* MH17-012N, *E. coli* MH13-051M (ST405), *K. pneumoniae* MH15-258M (ST11), and *K. pneumoniae* MH15-208H (ST395)] according to the following protocol. The same amount of Luria-Bertani (LB) broth cultures of each donor bacteria and the recipient azide– resistant *E. coli* J53 (ATCC BAA-2731, *F- met pro Azi^r^*), were mixed and spotted onto Mueller-Hinton agar and then incubated at 37°C overnight. The mixed cells were recovered and suspended into PBS buffer, plated onto LB agar after 10-fold serial dilution, and incubated at 37°C overnight. Transconjugants were selected on LB agar containing 2 μg/mL of meropenem and 100 μg/mL of sodium azide.

## Acknowledgements

We thank Meiko Takeshita for technical assistance and are grateful to the participating medical institution for providing the bacterial isolates and clinical information.

---

**Table S1.** Bacterial species and isolates, specimen types for the bacterial isolation, years in which the bacteria were isolated, and minimum inhibitory concentrations (MICs) of meropenem (MEM) and imipenem (IPM) of isolates are shown. Also, sequence types of multilocus sequence typing (MLST) analysis determined from genomes, carbapenemase genes detected by ResFinder in genomes, sizes and contigs of genomes, coverages in short-read sequencing, Illumina sequencing platforms, and accession numbers of genomes are shown. According to the CLSI 2020 guidelines, Breakpoints of meropenem and imipenem are as follows: ≤1 μg/mL, susceptible; 2 μg/mL, intermediate; ≥4 μg/mL, resistant (R).

| Species | Isolate | Specimen | Year | MEM MIC | IPM MIC | MLST | Carbapenemase gene | Size | Contig | Coverage | Platform | Accession no. |
| --- | --- | --- | --- | --- | --- | --- | --- | --- | --- | --- | --- | --- |
| *M. morganii* | MH16-367M | blood | 2016 | 1.5 | 8 (R) | N/A | *bla*NDM-1 | 4,132,245 bp | 297 | 88.2x | MiniSeq | BFCJ000000000 |
| *P. mirabilis* | MH13-009N | urine | 2013 | >32 (R) | >32 (R) | N/A | *bla*NDM-1 | 4,116,783 bp | 868 | 296.9x | MiniSeq | BFCK010000000 |
| *C. freundii* | MH17-012N | urine | 2017 | 2 | 3 | N/A | *bla*NDM-1 | 5,555,817 bp | 156 | 235.1x | HiSeq 4000 | BFCL010000000 |
| *C. freundii* | MH16-522D | pus | 2016 | >32 (R) | 12 (R) | N/A | *bla*NDM-1 | 5,311,537 bp | 182 | 119.4x | MiniSeq | BFCM010000000 |
| *E. coli* | MH17-008M | blood | 2017 | 4 (R) | 6 (R) | 457 | *bla*KPC-2 | 5,406,936 bp | 606 | 129.6x | HiSeq 4000 | BFCN010000000 |
| *E. coli* | MH13-051M | blood | 2013 | 6 (R) | 3 | 405 | *bla*NDM-1 | 5,656,255 bp | 259 | 107.9x | MiniSeq | BFCO000000000 |
| *K. pneumoniae* | MH17-004D | pus | 2017 | 4 (R) | >32 (R) | 11 | *bla*KPC-2 | 5,694,352 bp | 280 | 108.3x | HiSeq 4000 | BFCP000000000 |
| *K. pneumoniae* | MH15-289M | blood | 2015 | >32 (R) | >32 (R) | 11 | *bla*NDM-1 | 5,771,413 bp | 159 | 158.8x | HiSeq 4000 | BFCQ000000000 |
| *K. pneumoniae* | MH16-398D | pus | 2016 | 8 (R) | 2 | 11 | *bla*NDM-1 | 5,640,361 bp | 263 | 87.3x | MiniSeq | BFCR000000000 |
| *K. pneumoniae* | MH15-258M | blood | 2015 | 6 (R) | 2 | 11 | *bla*NDM-1 | 5,868,091 bp | 377 | 444.1x | MiniSeq | BFCS000000000 |
| *K. pneumoniae* | MH15-208H | sputum | 2015 | >32 (R) | >32 (R) | 395 | *bla*NDM-1 | 5,685,569 bp | 202 | 216.5x | HiSeq 4000 | BFCT000000000 |
| *K. pneumoniae* | MH15-191M | blood | 2015 | >32 (R) | >32 (R) | 395 | *bla*NDM-1 | 5,787,568 bp | 375 | 350.8x | MiniSeq | BFCU010000000 |
| *K. pneumoniae* | MH13-055M | blood | 2013 | >32 (R) | >32 (R) | 395 | *bla*NDM-1 | 5,718,215 bp | 248 | 255.0x | HiSeq 4000 | BFCV000000000 |
| *K. pneumoniae* | MH17-010D | pus | 2017 | >32 (R) | >32 (R) | 16 | *bla*NDM-4, *bla*OXA-181 | 5,608,215 bp | 160 | 261.8x | HiSeq 4000 | BFCW010000000 |
| *K. pneumoniae* | MH17-007D | pus | 2017 | 2 | 3 | 656 | *bla*KPC-2 | 5,637,531 bp | 149 | 131.1x | HiSeq 4000 | BFCX000000000 |
| *K. pneumoniae* | MH16-390M | blood | 2016 | 8 (R) | 8 (R) | 656 | *bla*NDM-1 | 5,638,714 bp | 219 | 97.1x | MiniSeq | BFCY000000000 |
| *K. pneumoniae* | MH17-556D | pus | 2017 | >32 (R) | >32 (R) | 15 | *bla*NDM-4 | 5,533,931 bp | 132 | 126.3x | HiSeq 4000 | BFCZ010000000 |
| *K. pneumoniae* | MH16-335M | blood | 2016 | 16 (R) | 16 (R) | 15 | *bla*NDM-4 | 5,840,408 bp | 274 | 140.2x | MiniSeq | BFDA000000000 |
| *K. pneumoniae* | MH17-016D | pus | 2017 | >32 (R) | 8 (R) | 15 | *bla*NDM-4, *bla*OXA-48 | 5,871,712 bp | 429 | 64.8x | MiniSeq | BFAI010000000 |
| *K. pneumoniae* | MH17-017D | pus | 2017 | 6 (R) | 8 (R) | 15 | *bla*NDM-4 | 5,847,449 bp | 250 | 203.0x | HiSeq 4000 | BFDB000000000 |
| *K. pneumoniae* | MH16-444M | blood | 2016 | 6 (R) | 4 (R) | 15 | *bla*KPC-2 | 5,640,233 bp | 288 | 122.7x | MiniSeq | BFDC000000000 |
| *K. pneumoniae* | MH17-553M | blood | 2017 | >32 (R) | >32 (R) | 15 | *bla*KPC-2 | 5,673,369 bp | 177 | 138.2x | HiSeq 4000 | BFDD010000000 |
| *K. pneumoniae* | MH17-536D | pus | 2017 | 3 | 8 (R) | 15 | *bla*KPC-2 | 5,652,240 bp | 292 | 74.8x | MiniSeq | BFDE000000000 |
| *K. pneumoniae* | MH16-519D | pus | 2016 | 6 (R) | 12 (R) | 15 | *bla*KPC-2 | 5,682,307 bp | 287 | 78.7x | MiniSeq | BFDF000000000 |
| *K. pneumoniae* | MH17-011M | blood | 2017 | >32 (R) | >32 (R) | 15 | *bla*KPC-2 | 5,652,498 bp | 202 | 182.1x | HiSeq 4000 | BFDG000000000 |

**Table S2.** Bacterial species and isolates, and *bla*_NDM_-carrying plasmids are shown. Also, replicon types detected by PlasmidFinder and antimicrobial resistance genes (ARGs) detected by ResFinder in plasmids, sizes of plasmids, and coverages in long-read sequencing, and accession numbers of plasmids are shown.

| Species | Isolate | Plasmid | Replicon type | ARG<br>(β-lactam) | ARG<br>(aminoglycoside) | ARG<br>(quinolone) | ARG<br>(others) | Size | Coverage | Accession no. |
| --- | --- | --- | --- | --- | --- | --- | --- | --- | --- | --- |
| *M. morganii* | MH16-367M | pMH16-367M_1 | IncFII(pSE11)<br>::IncN | *bla*NDM-1, *bla*TEM-1B | *rmtB* | *qepA1* | *tet(A)* | 73.1 kb | 1860.7x | AP018565 |
| *P. mirabilis* | MH13-009N | pMH13-009N_1 | IncFII(pSE11)<br>::IncN | *bla*NDM-1, *bla*TEM-1B | *rmtB* | *qepA1* | *tet(A)* | 72.6 kb | 758.8x | AP018566 |
| *C. freundii* | MH17-012N | pMH17-012N_1 | IncR | *bla*NDM-1, *bla*TEM-1B | N.D. | N.D. | N.D. | 39.2 kb | 930.6x | AP018567 |
| *C. freundii* | MH17-012N | pMH17-012N_2 | IncFII(pHN7A8)<br>::IncN | N.D. | N.D. | N.D. | N.D. | 87.6 kb | 930.6x | AP018568 |
| *C. freundii* | MH16-522D | pMH16-522D_1 | IncX3 | *bla*NDM-1, *bla*SHV-12 | N.D. | N.D. | N.D. | 52.7 kb | 924.5x | AP018571 |
| *E. coli* | MH13-051M | pMH13-051M_1 | IncFII(pRSB107)<br>::IncN | *bla*NDM-1, *bla*CTX-M-55,<br>*bla*TEM-1B | *rmtB* | N.D. | *fosA3* | 111.5 kb | 388.7x | AP018572 |
| *K. pneumoniae* | MH15-289M | pMH15-289M_1 | IncFIA(HI1)<br>::IncA/C2 | *bla*NDM-1, *bla*CTX-M-15,<br>*bla*OXA-1 | *rmtC*, *aac*(6')-Ib-cr,<br>*aac*(3)-IIa, *aac*(3)-IId | *qnrB9* | *catA2*, *catB3*, *sul1*,<br>*dfrA14* | 147.8 kb | 125.0x | AP018577 |
| *K. pneumoniae* | MH16-398D | pMH16-398D_1 | IncFIA(HI1)<br>::IncA/C2 | *bla*NDM-1, *bla*CTX-M-15,<br>*bla*OXA-1 | *rmtC*, *aac*(6')-Ib-cr,<br>*aac*(3)-IIa, *aac*(3)-IId | *qnrB9* | *catA2*, *catB3*, *sul1*,<br>*dfrA14* | 147.7 kb | 224.1x | AP018578 |
| *K. pneumoniae* | MH15-258M | pMH15-258M_1 | IncFIA(HI1)<br>::IncA/C2 | *bla*NDM-1, *bla*CTX-M-15,<br>*bla*OXA-1 | *rmtC*, *aac*(6')-Ib-cr,<br>*aac*(3)-IIa, *aac*(3)-IId | *qnrB9* | *catA2*, *catB3*, *sul1*,<br>*dfrA14* | 148.8 kb | 584.9x | AP018579 |
| *K. pneumoniae* | MH15-208H | pMH15-208H_1 | IncFII(Yp) | *bla*NDM-1 | *rmtB* | N.D. | N.D. | 75.4 kb | 252.3x | AP018580 |
| *K. pneumoniae* | MH15-191M | pMH15-191M_1 | IncFII(Yp) | *bla*NDM-1 | *rmtB* | N.D. | N.D. | 75.6 kb | 788.4x | AP018581 |
| *K. pneumoniae* | MH13-055M | pMH13-055M_1 | IncFII(Yp) | *bla*NDM-1 | *rmtB* | N.D. | N.D. | 75.3 kb | 243.8x | AP018582 |
| *K. pneumoniae* | MH16-390M | pMH16-390M_1 | IncFIA(HI1)<br>::IncR | *bla*NDM-1, *bla*CTX-M-3,<br>*bla*TEM-1B | *aac*(6')-Ib-cr, *aadA16* | *qnrB6* | *ARR-3*, *sul1*, *dfrA27* | 345.7 kb | 431.1x | AP018583 |
| *K. pneumoniae* | MH16-335M | pMH16-335M_1 | IncFII(K) | *bla*NDM-4, *bla*OXA-9,<br>*bla*SHV-106, *bla*SHV-28 | *aac*(6')-Ib-cr, *aadA1* | N.D. | N.D. | 105.6 kb | 180.6x | AP018584 |

